# Population coupling of V1 and V4 neurons and its relation to local cortical state fluctuations and attention in macaque monkey

**DOI:** 10.1101/2025.09.19.677049

**Authors:** Mahyar Doost, Jochem van Kempen, Michael Boyd, Alexander Thiele

## Abstract

Neurons couple to various degrees to the activity level of the local neighboring population whereby strongly coupled ‘choristers’ and weakly coupled ‘soloists’ have been identified as two extremes of a continuous spectrum. At the same time neuronal populations undergo coordinated ON and OFF cortical state activity fluctuations, which are locally modulated by attention. The population coupling of soloists and choristers suggests that soloists should show limited alignment with cortical state fluctuations, while choristers should exhibit profound alignment. To test this, we recorded neurons across cortical layers in macaque areas V1 and V4, while animals performed a feature based spatial attention task. As expected, we found a wide range of population coupling strength of neurons. In line with our prediction, coupling of choristers to cortical state changes (ON-OFF transitions) was generally stronger than that of soloists. The strength of population coupling of neurons was similar during spontaneous and stimulus driven activity. Allocation of attention to the receptive field reduced the population coupling strength. Attentional modulation of neurons was positively correlated with population coupling strength. While neurons on average retained their coupling strengths across conditions, some neurons change coupling strength condition dependent, thereby potentially enhancing the coding abilities of cortical circuits.

## Introduction

Global brain states manifest as large-scale spatiotemporal activity pattern ^1,2^. They evolve on multiple temporal scales, range from minutes to hours during different sleep stages ^3^, and from seconds to minutes during wakefulness ^4-6^. Brain states affect neuronal responses, neural interactions, and neural information encoding ^4,6-9^. They underlie arousal and link to cognitive function and conscious perception ^6,9^. Cortical state fluctuations manifest in single cortical columns by spontaneously transitioning between states of vigorous (On) and faint (OFF) spiking periods ^10-12^. Initially, it was assumed that these fluctuations affect activity homogenously throughout the cortex ^4,13^. However, recent studies show that endogenous fluctuation of cortical states are locally modulated by attention, and the local momentary state (ON or OFF) predicts detection of visual stimuli and reaction times ^10,11^.

The alternation of population activity between ON-OFF spiking periods suggests that neurons behave in unison. However, various studies have shown that neurons have distinct activity pattern and their coupling strength with the neighboring neural population can vary radically. The coupling strength can be quantified as population coupling (PC) ^14^. Strongly coupled ‘choristers’ and weakly coupled ‘soloists’ have been identified as two extremes of a continuous spectrum ^12,14-17^. This raises the question, to what extent soloists exhibit ON-OFF spiking activity and whether soloists have a more continuous mode of functioning during ON-OFF fluctuations when compared to choristers. Given the local modulation of cortical state with attention it also raises the question whether soloists and choristers show different levels of attentional modulation. Flexibility in coupling strength might be an important component of increasing coding abilities. Soloists may have the ability to flexibly join multiple different neural ensembles to perform condition dependent encoding. If such flexibility was also permitted to some choristers, it would further expand the encoding potential of the circuit ^18^. To investigate whether condition dependent flexibility of coupling strength is present in soloist and choristers, we recorded from laminar probes across cortical layers simultaneously in macaque area V1 and V4, while monkeys engaged in a feature based spatial attention task (figure 1). We quantified how soloists and choristers couple to ON-OFF period fluctuations and examined whether PC of soloists and choristers was differently affected by spatial attention. We found that PC was largely a fixed feature across V1 and V4 populations, but coupling strength in some neurons showed condition dependent variation.

**Figure 1:**
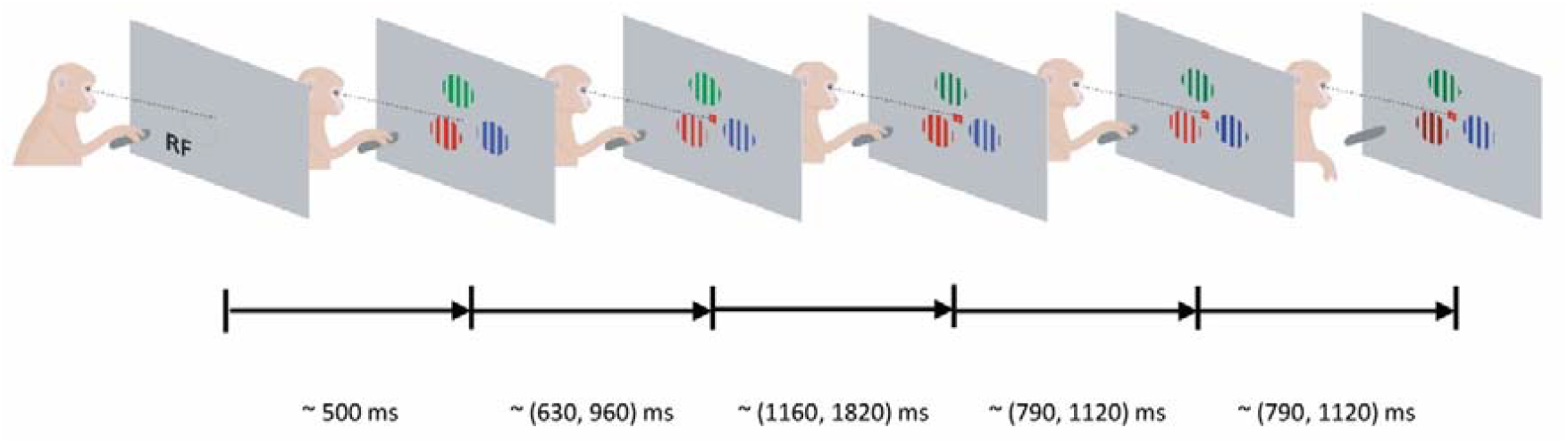
Feature-based spatial attention task. The monkey held a lever to initiate the trial, whereafter a central fixation spot was turned on. Upon fixation 3 colored gratings appeared, one was centered on the receptive fields (RFs) of the V1 neurons. After a variable delay a cue matching one of the grating colors was presented centrally along with the fixation spot, indicating which grating was behaviorally relevant (target). In pseudorandom order the stimuli decreased in luminance (dimmed). Upon dimming of the target, the monkey had to release the lever to receive a reward.

## Results

### Population coupling

Monkeys performed a cued feature based spatial attention task (figure 1, Methods). We first describe basic aspects of PC of individual neurons during different epochs of the task, and different attentional conditions. In line with previous reports by Okun and colleagues in 2015 ^14^ we found that PC varied widely between individual neurons (Figure 2B-E), and it varied between cortical layers, being generally strongest in supragranular layers (Supplementary Figure S1).

**Figure 2:**
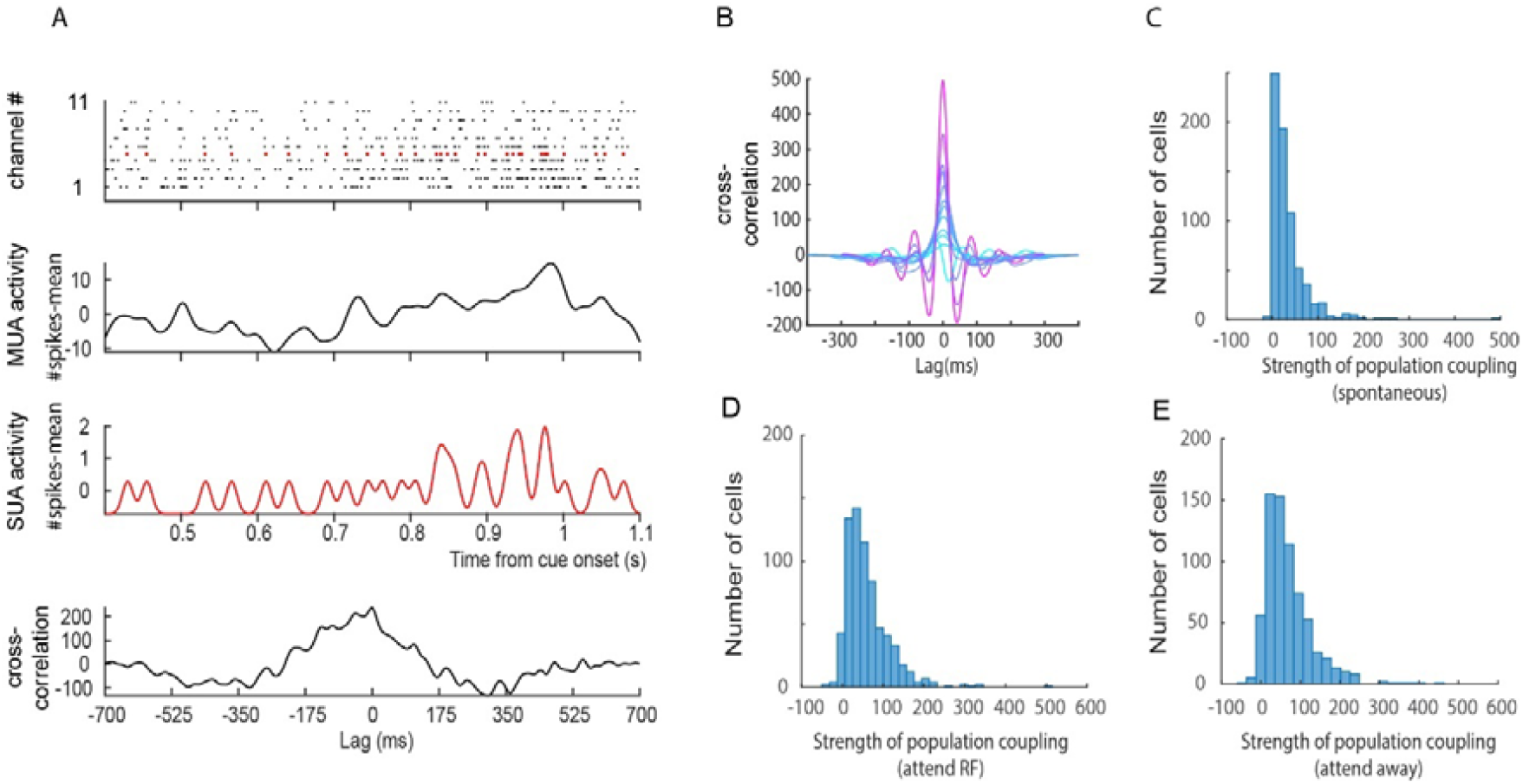
Calculation of population coupling and its distribution for different experimental epochs and conditions. **A)** Raster plots within a single trial of MUA activity (black) across different channels located in grey matter. Red raster plot shows SUA from a channel, recorded along the MUA activity. Averaged, smoothed and mean subtracted MUA activity across channels is shown below raster plots. Smoothed and mean subtracted SUA activity is shown in red. Bottom plot in A shows cross correlation between MUA activity and SUA activity for this trial. **B)** Range of population coupling shown for selected cells. Coupling strength was determined from the height of the cross-correlogram at 0ms time lag. Color coding shows coupling from least coupled example cells (light blue) to most coupled cells (magenta). **C)** Distribution of coupling strength during spontaneous activity. **D)** Distribution of coupling strength after cue onset during attend RF conditions. **E)** Distribution of coupling strength after cue onset during attend away conditions.

We next investigated whether PC varied for different stimulus conditions. Here we compared PC during spontaneous activity and during stimulus driven activity. Figure 2 C-E shows data pooled across areas. At first glance coupling strength seems similar during spontaneous activity and post cue onset activity, but they differed significantly between areas and periods (details see table 2), which we delineate in the next sections. To further investigate to what extent coupling strength is a largely fixed feature of a neuron, we calculated the correlation between coupling strength during spontaneous activity and after cue onset. We used the normalized population coupling of SUAs for each condition (C_norm_ and its Fisher Z-transform Z(0), see Methods). The strength of population coupling of SUAs during spontaneous and stimulus driven activity was highly correlated for both areas (Figure 3A-D), and the correlations were similar and significant for each monkey. This demonstrates that under these conditions the PC is largely a fixed feature of a neuron. As described in Methods, we additionally controlled for the possibly that single-unit activity leaked into neighboring MUA channels, or that neurons would not respond to the stimulus, which both might affect the results reported. Neither was the case as shown in Supplementary Figures S2.

**Figure 3:**
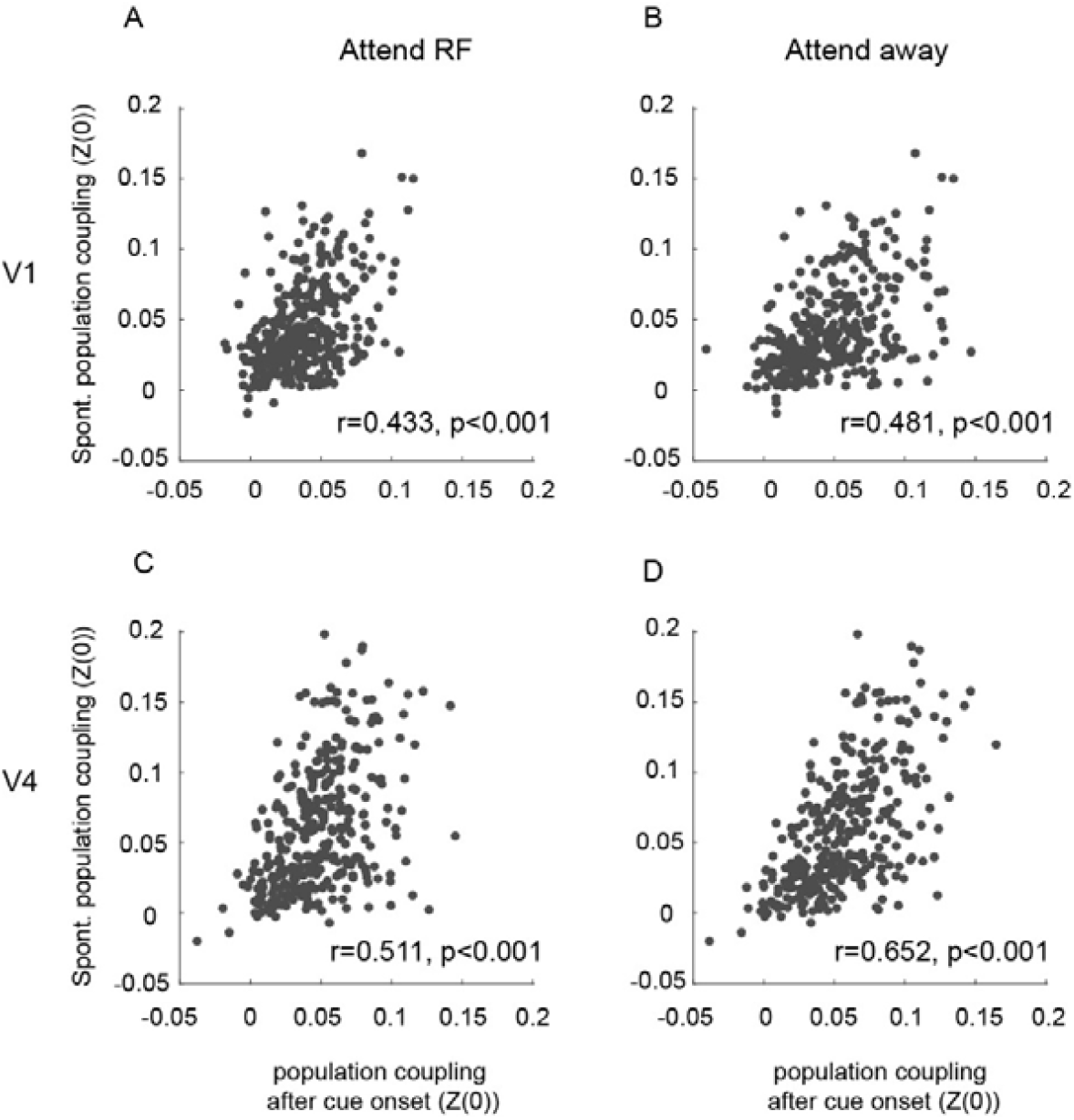
Population coupling during spontaneous activity compared to stimulus driven activity. **A)** Correlation of Population coupling during spontaneous activity and during attend RF stimulus driven activity for V1 neurons (period after cue onset). **B)** Correlation of Population coupling during spontaneous activity and during attend away stimulus driven activity for V1 neurons (period after cue onset). **C)** Correlation of Population coupling during spontaneous activity and during attend RF stimulus driven activity for V4 neurons (period after cue onset). **E)** Correlation of Population coupling during spontaneous activity and during attend away stimulus driven activity for V4 neurons (period after cue onset). Insets show correlation coefficient between the two measures, and respective p-values. Population coupling was quantified by calculating the Fisher transformed correlation coefficient between single unit and population activity (Z(0)).

Despite this pattern, PC may still be modulated by cognitive factors such as attention or cortical state fluctuations, which we explore below.

### Effect of spatial attention on population coupling

Previously Okun et al. ^14^ showed that behavioral variables such as motor intention influenced population coupling of V4 neurons. Here we evaluate the effect of covert spatial attention on the strength of population coupling of neurons. To compare the strength of population coupling of neurons between attend RF and attend-away conditions, we used the Fisher transformed PC correlation coefficient of a neuron (Z(0), see Methods), and compared these between attention conditions. Figure 4 shows the scatter plot of normalized PCs for attend RF vs attend away conditions for areas V1 and V4, respectively. The majority of data points lie above the diagonal, and consequently PC was significantly stronger during attend out conditions compared to attend RF conditions in both areas (p<0.001, Wilcoxon signed rank test, results were significant for each monkey at p<0.001). This demonstrates that allocation of attention to the RF decreased population coupling of neurons (For controls relating to ‘leakage’, rate differences, and stimulus responsiveness between attention conditions see Supplementary Figures S3). Attentional modulation itself differed between layers in V1, it was weakest in granular layers (Supplementary Figure S4), but no significant differences were found in V4 (Supplementary Figure S4).

**Figure 4:**
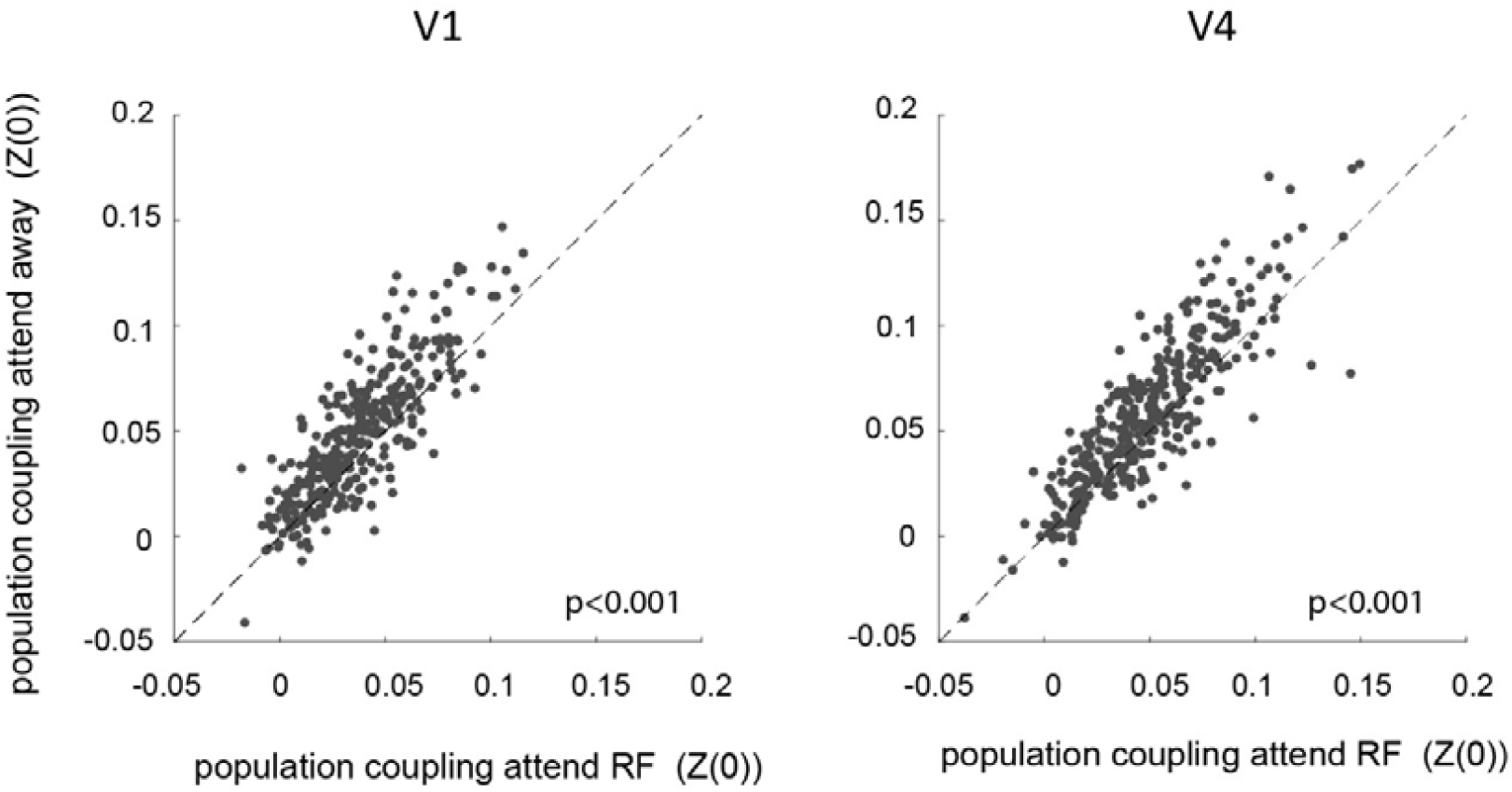
Effect of spatial attention on population coupling of V1 and V4 neurons. Population coupling (Fisher-transformed correlation coefficient (Z(0)) during attend RF conditions (x-axis) and attend away conditions (y-axis) during the period after cue-onset for V1 neurons (left) and V4 neurons (right). Insets show p-values of differences in population coupling.

### Coupling of soloists and choristers to population state change transitions

Recent studies have demonstrated that attention locally affects brain state. Brain state manifests as ON-OFF transitions of activity, whereby attention to the receptive field increases ON-period durations in V1 and V4, in addition to increasing neural activity during both ON- and OFF-periods ^10,11^. Given the broad range of PCs across neurons, we would expect that soloists follow ON-OFF transitions to a lower degree than choristers. Note, that while this prediction is straightforward, it is not a tautology, as the PC calculations have been performed without regard of ON-OFF period activity itself, and it has not been tested explicitly before. To examine the coupling of soloists and choristers with cortical state fluctuations during stimulus driven activity, we first calculated the ratio of firing rates during ON to OFF phases for each neuron. This provides a measure how much a neuron changes its activity with ON-OFF transitions. We then calculate the correlation of this ratio with the PC (Z(0)) of the neurons. The results are shown in (**Figure 5**, see Supplementary Figure S5 for leakage and responsiveness controls, respectively). As predicted, on average cells with lower population coupling, also change their activity rates less during ON-OFF phase transitions during either attention condition, which is evident from the positive correlation between PCs versus ratio of firing rates during ON to OFF phases (p<0.001 for both areas and attention conditions). However, the correlation coefficients themselves only explain between ∼10% to ∼24% of the variance, and figure 5 demonstrates that some soloists (PC close to 0) nevertheless show profound activity differences during ON to OFF phase transitions (ratios of 4 to 6).

**Figure 5:**
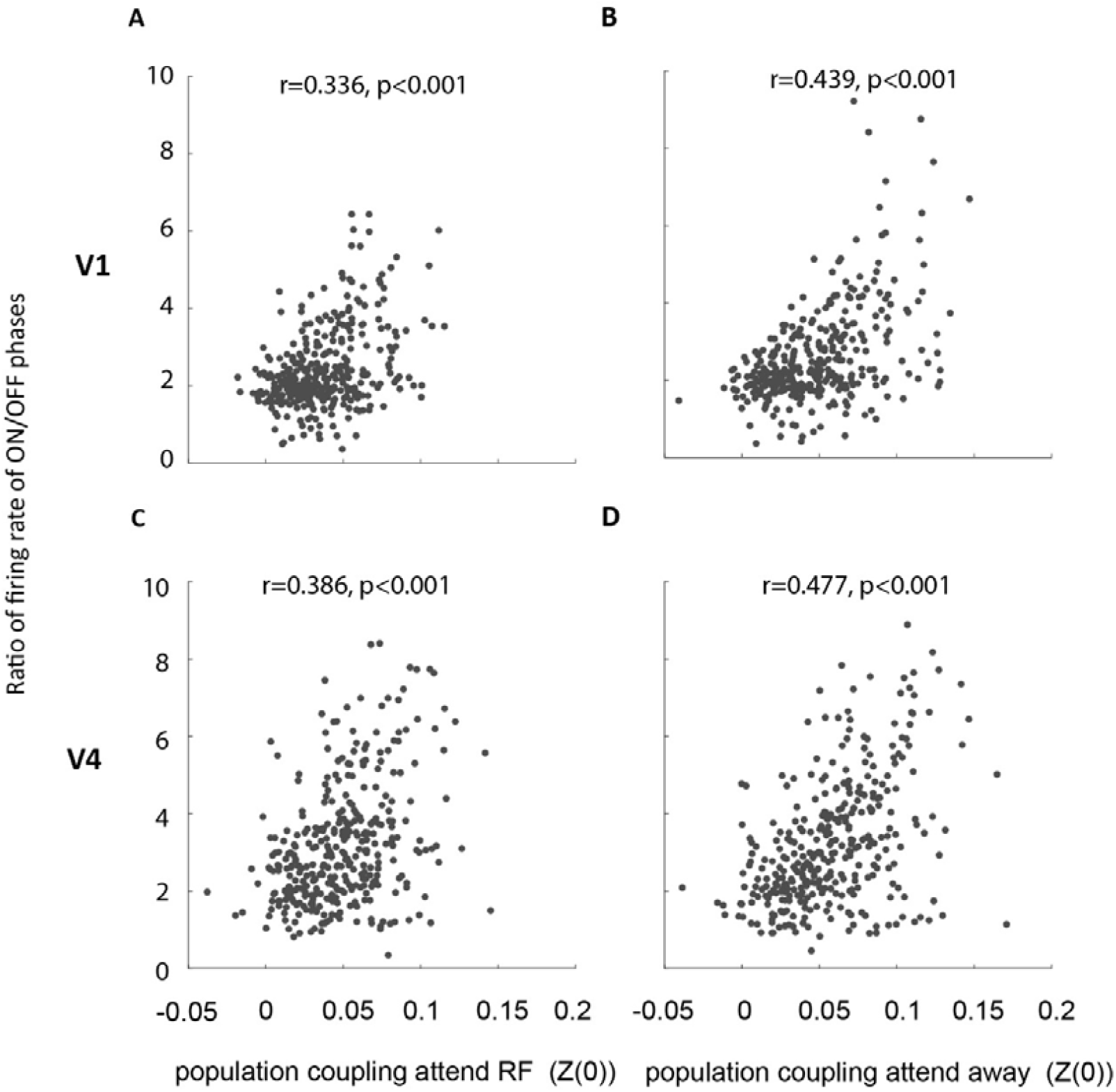
Relation between ON-OFF state fluctuation alignment and population coupling of V1 and V4 neurons. Population coupling (Fisher-transformed correlation coefficient Z(0)) during attend RF conditions and attend away conditions during the period after cue-onset for V1 neurons (left) and V4 neurons (right) compared to the ratio of ON-OFF period firing (y-axis). Insets show correlation coefficients and p-values of this relationship.

### Attentional modulation of soloists and choristers

Given that soloists and choristers (on average) differentially align with cortical state ON - OFF phase transitions, which themselves are affected by spatial attention, we asked whether soloists and choristers are differently affected by spatial attention. To this end we calculated an attentional modulation index for each neuron (Methods) and correlated this with the PC of the neurons (**Figure 6**, also see Supplementary Figure S6 for ‘leakage’ and responsiveness controls). A positive correlation would indicate that choristers show stronger attentional modulation, while a negative correlation would indicate the opposite. We find that in both V1 and V4 there is a positive correlation between the attentional modulation index and the PC strength (V1: r= 0.158, p=0.003, V4: r= 0.199, p<0.001). Thus, there is a tendency for choristers to exhibit stronger attentional modulation when compared to soloists, but overall, this tendency is modest (the correlation explains 2.5-4% of the variance).

**Figure 6:**
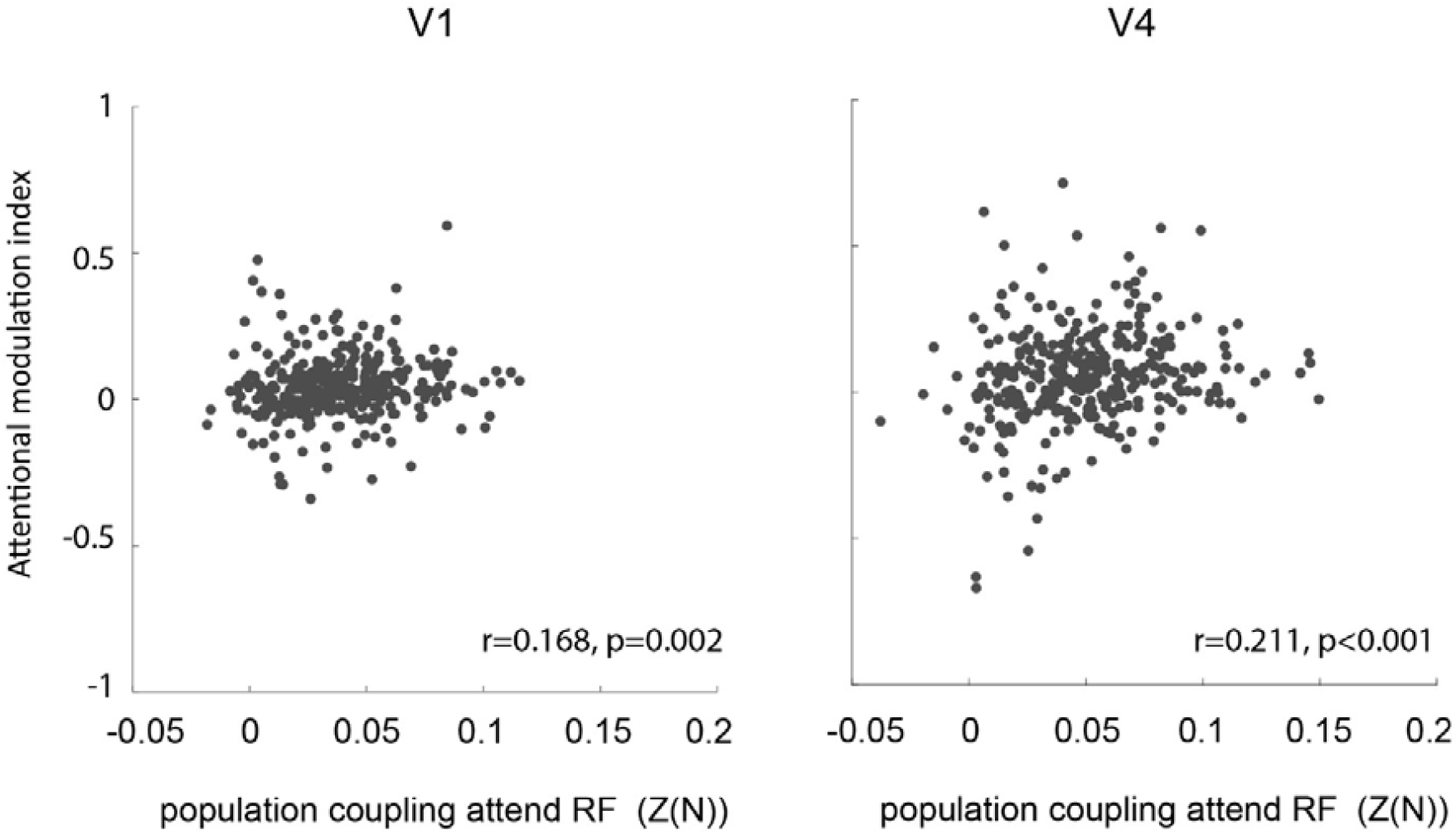
Relation between population coupling of V1 and V4 neurons and their attentional modulation strength. Population coupling (Fisher-transformed correlation coefficient (Z(0)) during attend RF conditions during the period after cue-onset for V1 neurons (left) and V4 neurons (right) compared to the strength of attentional modulation (y-axis). Insets show correlation coefficients and p-values of this relationship.

### Relationship between population coupling and noise correlation

Population coupling should be related to noise correlation measures, as both assess co-variability of neural firing. While the former does so on a larger scale (one neuron’s coupling to the local population activity), the latter assesses single neuron pairwise coupling, usually over somewhat longer time scales (often >100 ms integration windows). We did find that attention reduced noise correlations in V1, and in V4 (supplementary figure S7A). This is in line with the effects of attention on population coupling. We also predicted that neurons with strong population coupling (on the chorister spectrum end), would show stronger pairwise coupling (larger pairwise noise correlations). We found this to be true as well (supplementary figure S7B), but overall the effect, while significant, was small, explaining ∼5-8% of the variance.

## Discussion

We show that the strength of population coupling of neurons varied widely in macaque area V1 and V4. In general, the strength of population coupling was similar during spontaneous and stimulus driven activity. Population coupling during attend RF condition was smaller than during attend away condition. Choristers coupled more strongly with cortical state changes (ON-OFF transitions) than soloists and showed stronger attentional modulation. Despite these general tendencies we also found that PC can vary in a condition dependent manner for some neurons.

In line with previous studies we find that PC of neurons forms a continuous spectrum ranging from strongly coupled ‘choristers’ to weakly coupled ‘soloists’ ^12,14-17^. These can be identified in both areas V1 and V4 of the macaque, whereby PC strength was strongly correlated between spontaneous and stimulus driven activity, as well as during different attention conditions. However, PC was generally stronger during attend away conditions when compared to attend RF conditions, i.e. PC is modulated by cognitive factors. This is in line with previous reports, where motor intention affected PC strength of V4 neurons ^14^.

Cortical state fluctuations manifest in single cortical columns by spontaneously transitioning between states of vigorous (On) and faint (OFF) spiking periods ^10-12^. These transitions are locally affected by spatial attention, and we found that soloists generally couple less to the ON-OFF phase activity transitions when compared to choristers, as could have been predicted by the underlying population coupling behavior. However, it is important to note that there are exceptions to the rule, whereby neurons with close to zero PC values still show profound activity changes during ON-OFF transitions, and neurons with strong PC values, exhibit very limited activity changes during ON-OFF transitions (see some individual data points in figure 5).

How does population coupling relate to more traditional measures such as noise correlations? Noise correlations only measure pairwise coupling, while population coupling is a more large scale measure. We found that noise correlations were reduced by attention in line with numerous previous reports ^19-25^, and we equally found that noise correlations between pairs of neurons was significantly correlated with their individual population coupling strength. However this correlation, while significant, was relatively small, explaining less 5-8% of the variance. This is not entirely surprising, given that noise correlations between neurons can be positive or negative, whereby the population coupling would be affected by all pairwise combinations (positive and negative), which could thus average to zero even if a single pair shows a large correlation.

Given the local modulation of cortical state with attention, and the stronger coupling of choristers with ON-OFF state transitions we predicted that choristers should show stronger attentional modulation when compared to soloists. This prediction turned out to be correct, but the overall alignment was limited, i.e. PC, while a significant predictor of attentional modulation, only explained a limited amount of variance of attentional modulation. It suggests that attention induced firing rate changes depend to a very small extent on population coupling and are thus possibly driven by a different mechanism.

Our data were recorded with laminar electrodes, which most likely sampled from single cortical columns, where coupling strength is more profound than between distant columns ^26-28^. To what extent our results map to population coupling and its modulation by attention between different columns remains to be determined.

In summary we found that PC of macaque V1 and V4 neurons on average shows limited variation across different stimulus periods, while it is affected by cognitive factors such as spatial attention. This may allow some neurons to flexibly engage in multiple different neural ensembles, which would greatly increase the coding capacity of a given neural population ^18^.

## Disclosures and acknowledgments

AT was funded by the MRC (MR/P013031/1), BBSRC (BB/W006758/1), *and by In2PrimateBrains (European Union’s Horizon 2020 research and innovation program under the Marie Skłodowska-Curie Actions Doctoral Networks, Grant Agreement No 956669)*.

## Data availability statement

The data relevant for the manuscript have been uploaded to https://gin.g-node.org/Mahyar-Doost/md-soloists-and-choristers-data.git. and will be available upon manuscript publication. All relevant analysis scripts have been deposited at https://gitlab.com/nmd174/md-soloists-and-choristers.git, which will be made publicly available upon manuscript publication.

## Materials and Methods

### Data set

The data set used had previously been recorded, consisting of data that contributed to different publications from our group ^11,29,30^.

### Subjects and general procedures

Two awake male monkeys (Macaca mulatta, age 10-12 years, weight 8-9.5 kg) were used as subjects in this study. They were housed under conditions described in detail previously ^31^. All procedures were approved by the Newcastle University Animal Welfare Ethical Review Board (AWERB) and the Animal in Sciences Committee UK, and carried out in accordance with the European Communities Council Directive RL 2010/63/EC, the US National Institutes of Health Guidelines for the Care and Use of Animals for Experimental Procedures, and the UK Animals Scientific Procedures Act.

### Surgical procedures

The animals were implanted with a head post and recording chambers over areas V1 and V4 under sterile conditions and general anesthesia. Surgical procedures and postoperative care conditions have been described in detail previously ^32^.

### Behavioral paradigm

The behavioral paradigm (**Figure 1**) has been described previously ^11,29^, but we copy the description here for ease of access.

Stimulus presentation and behavioral control was managed by Remote Cortex 5.95 (Laboratory of Neuropsychology, National Institute for Mental Health, Bethesda, MD). Stimuli were presented on a cathode ray tube (CRT) monitor at 120 Hz, 1280 × 1024 pixels, at a distance of 54 cm.

During the main task (**Figure 1**), the monkeys initiated a trial by holding a lever and fixating on a central white fixation spot (0.1°) displayed on a grey background (1.41 cd/m^2^). After a fixed delay (614, 424 ms for monkey 1 and 2), three colored square wave gratings (for color values see **Table 1**) appeared equidistant from the fixation spot, one was centered on the RF of the V1 neurons under study.

**Table 1.**
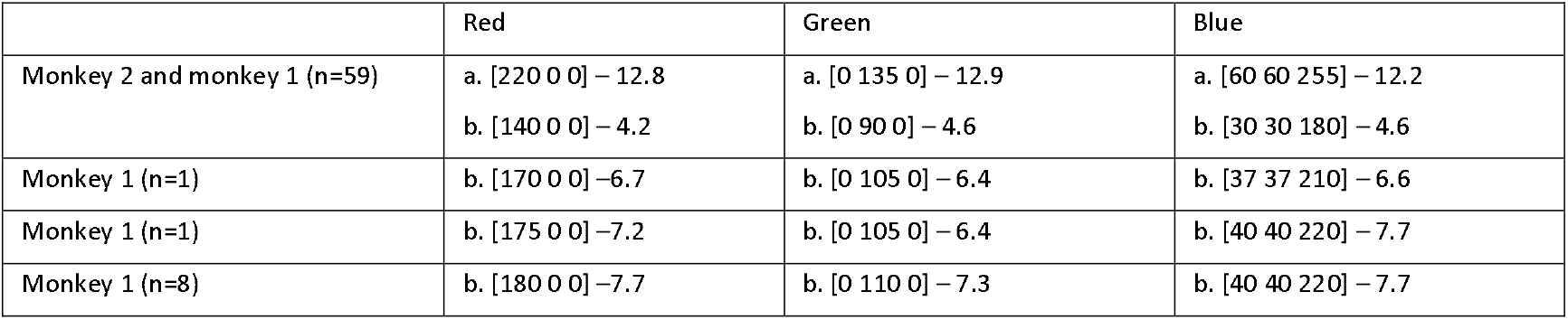
Color values used for the three colored gratings across recording sessions and subjects Methods. Color values are indicated as [RGB] – luminance (cd/m2). a = Undimmed values, b = Dimmed values. For monkey 1, we used a variety of dimmed values across recordings.

|  | Red | Green | Blue |
| --- | --- | --- | --- |
| Monkey 2 and monkey 1 (n=59) | a. [220 0 0] – 12.8<br>b. [140 0 0] – 4.2 | a. [0 135 0] – 12.9<br>b. [0 90 0] – 4.6 | a. [60 60 255] – 12.2<br>b. [30 30 180] – 4.6 |
| Monkey 1 (n=1) | b. [170 0 0] – 6.7 | b. [0 105 0] – 6.4 | b. [37 37 210] – 6.6 |
| Monkey 1 (n=1) | b. [175 0 0] – 7.2 | b. [0 105 0] – 6.4 | b. [40 40 220] – 7.7 |
| Monkey 1 (n=8) | b. [180 0 0] – 7.7 | b. [0 110 0] – 7.3 | b. [40 40 220] – 7.7 |

**Table 2.** Average, standard deviation and medians of population coupling (PC) in V1 and V4 across different time windows/conditions. P: p-value based on Wilcoxon rank sum test comparing PC between area V1 and V4. Lower rows shows p-values for comparisons across different periods within each area (Kruskal Wallis Anova with rank sum test post-hoc comparison).

|  | Spontaneous activity |  |  |  | Attend RF |  |  |  | Attend away |  |  |  |
| --- | --- | --- | --- | --- | --- | --- | --- | --- | --- | --- | --- | --- |
|  | Mean<br>PC | Std | Median | p | Mean<br>PC | Std | Median | p | Mean<br>PC | Std | Median | p |
| V1 | 24.75 | ±23.17 | 17.04 | <0.001 | 48.22 | ±39.65 | 38.64 | <0.001 | 58.52 | ±48.04 | 48.13 | <0.001 |
| V4 | 42.91 | ±50.61 | 28.62 |  | 72.07 | ±62.65 | 55.31 |  | 81.13 | ±74.34 | 63.21 |  |
|  | Difference between<br>periods (p) |  |  | p<sub>spont.-attend RF</sub> |  |  | p<sub>spont.-attend away</sub> |  |  | p<sub>attend RF-attend away</sub> |  |  |
| V1 | <0.001 |  |  | <0.001 |  |  | p < 0.05 |  |  | <0.001 |  |  |
| V4 | <0.001 |  |  | <0.001 |  |  | p = 0.3126 |  |  | <0.001 |  |  |

The locations of colored gratings were fixed for each recording session but were pseudo-randomly varied across sessions. Stimulus size varied between 2 and 4° diameter, depending on RF eccentricity and size. For most recordings we used drifting gratings but presented one monkey (monkey 2) with stationary gratings during 22 out of 34 recording days. The drifting gratings moved perpendicular to the grating orientation, with the motion direction pseudo-randomly assigned on every trial. After a random delay (618-1131 ms for monkey 1, 618-948 ms for monkey 2; uniformly distributed), a central cue appeared that matched the color of one of the gratings, indicating that this grating would be behaviorally relevant on the current trial. After a variable delay (1162-2133 ms for monkey 1, 1162-1822 ms for monkey 2; uniformly distributed), one of the gratings changed luminance (for color values see Table 1), referred to as dimming. If the cued grating (target) dimmed, the monkey had to release the lever in order to obtain a reward. If, however, a non-cued grating (distractor) dimmed, the monkey had to ignore this and keep hold of the lever until the target dimmed on the second or third dimming event (each after another 792-1331 ms for monkey 1; 792-1164 ms for monkey 2; uniformly distributed).

### Data acquisition and analysis

We recorded across cortical layers of visual areas V1 and V4 using 16-contact laminar electrodes (150 µm contact spacing, Atlas silicon probes). We included recordings where the Hidden Markov model analysis (see below) suggested that a 2-phase model best approximated the data. This applied to 34 penetrations from monkey 1 and 35 from monkey 2. Of these, 28 penetrations were performed in V1 and 29 were performed in V4 of monkey 1 with 26 performed simultaneously in V1 and V4. In monkey 2 all 35 included penetrations were from simultaneous recordings in V1 and V4. The electrodes were inserted perpendicular to the cortex on a daily basis.

Raw data were collected using Remote Cortex 5.95 and by Cheetah (Neuralynx) data acquisition interlinked with Remote Cortex 5.95. Neuronal data were acquired with Neuralynx preamplifiers and a Neuralynx Digital Lynx amplifier. Unfiltered data were sampled with 24 bit at 32.7 kHz and stored to disc. Data were replayed offline, sampled with 16-bit and band-pass filtered at 0.5-300 Hz and down sampled to 1 kHz for local field potential (LFP) data, and filtered at 0.6-9 kHz for spike extraction sampled at 32.7 kHz. Eye position and pupil diameter was recorded at 220 Hz (ViewPoint, Arrington Research).

All data analyses were performed using custom written Matlab (the Mathworks) scripts.

### Data preprocessing

We corrected for any noise common to all channels via common average referencing, whereby the average of all channels of an electrode was subtracted from each individual channel of that electrode.

In order to determine signal-to-noise (SNR) ratios, recording stability, the visual response latency, as well as for the computation of receptive fields (i.e. for preprocessing purposes only), we computed the envelope of MUA (MUAe) by low-pass filtering (<300 Hz, fifth order Butterworth) the rectified 0.6-9 kHz filtered signal. During some recording sessions the electrode moved (e.g. due to movement of the monkey). To account for this, we visually inspected the stability of each recording by investigating the stimulus aligned firing rates, MUAe and their baseline (-500 to -50 ms) energy across all trials and channels. With energy (*E*) defined as:

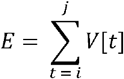

where *t* is the running index in the vector (*V*) representing the single-trial histogram (or MUAe), *i* is the first time point of the trial and *j* is the last time point of it. We selected the largest continuous time window that showed stable activity across all V1 and V4 channels.

Single unit activity (SUA) was extracted by band pass filtering the neural signal between 600 and 9,000 Hz, followed by manual spike sorting using SpikeSort3DSetup (Neuralynx) software. We included SUAs with at least 5Hz firing rate in any time window of interest (see below) and for which we recorded at least 10 trials per attention condition. Multi-unit spiking activity was extracted in the same way, but here the criteria of single unit isolation (less than 1% of entries in the first ms bin of the interspike interval) were not fulfilled.

### Layer compartments alignment

To identify recording channels located in the gray matter of V1 and V4, a current source density (CSD) analysis was carried out. Here current sinks and sources allowed to determine the relative recording depth, which was then compared to the known cortical anatomy ^33^. The CSD profile can be calculated according to the finite difference approximation, taking the inverse of the second spatial derivative of the stimulus-evoked voltage potential φ, defined by:

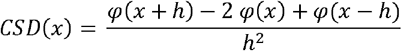

where *x* is the depth at which the CSD is calculated and *h* the electrode spacing (150 μm). We used the iCSD toolbox ^34^ to compute the CSD. With this toolbox we used a spline fitting method to interpolate φ smoothly between electrode contacts. We used a diameter of cortical columns of 500 µm ^35^, and tissue conductance of 0.4 Sm^-1 36^. The location of the early sink was identified visually by using interactive CSD plots, whereby ‘clicking’ on the location of the early sink saved the depth coordinates in relation to all electrode contacts. This in conjunction with the latency analysis (see below) allowed to align depths across sessions, and assign laminar location to a contact.

To aid determination of recording depth, we also computed the signal-to-noise ratio (SNR), the response latencies to stimulus onset for each channel and the receptive field (RF, see below). SNR was computed as:

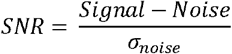

with *Singnal* defined as the average MUAe amplitude in one of eight 50 ms time windows, from 30 to 80 ms, in 10 ms steps, to 100 to 150 ms after stimulus onset, and *Noise* defined as the average MUAe amplitude during the baseline period (-200 to 50 ms) before stimulus onset. SNR in at least one of these eight estimates was required to be higher than 3 for a channel to be included for further analyses.

We computed the response latency to stimulus onset for each channel according to a method previously described ^37^. We fitted the visual response as a combination of an exponentially modified Gaussian and a cumulative Gaussian using a non-linear least-squares fitting procedure (function lsqcurvefit) applied to the average MUAe time course. There are two assumptions implicit in this method. First, the onset latency has a Gaussian distribution across trials and across neurons that contribute to the MUAe, and second, that (part of) the response dissipates exponentially. The visual response *y* across time *t* was modelled as:

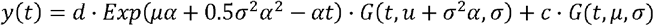

where μ is the mean, σ is the standard deviation, α^-1^ is the time constant of the dissipation, *G* (*t*, μ, σ )is a cumulative Gaussian, and *c* and *d* are the factors scaling the non-dissipating and dissipating modulation of the visual response. The response latency was defined as the time point where *y* (*t*) reached 33% of the maximum of the earliest peak of the Gaussian ^37^. The earliest current sink/earliest latency was taken as the presumed thalamic input layer (L4) reference channel. In area V1, channels located 0.25 to 1 mm above the reference channel were assigned to supragranular, channels located within +/-0.25 mm were assigned to granular, and channels located 0.25 to 0.75 mm below the reference channel were assigned to the infragranular layer compartment. It assumes a thickness of ∼0.5 mm of the granular layer which is in line with published results e.g. ^38,39^. In area V4, channels at distances of 0.1 to ∼1 mm above the reference channel were labeled as supragranular, channels above or below the reference channel within 0.1 mm were labeled as granular, and channels at distances of 0.1 to ∼0.75 mm below the reference channel were labeled as being in the infragranular layer compartment. This assumes a layer 4 thickness of ∼200 um, which estimates it to be thinner than layer 4 of V1 in line with the existing literature V4 layer thickness was estimated based on figures in previous publications e.g. ^40,41,42^. While such a thickness may be on the conservative side, it ensures that cells outside the granular layer are not misclassified as granular cells. Since the latter are a smaller sample, misclassification would have larger effects on comparative measures than misclassifying granular cells as supra- or infragranular. For assignment to supra- and infra-granular layers we also used the presence of spiking activity at a given channel. If a channel was >1mm above the granular layer in V1, but showed clear spiking activity we still assigned it to be in supragranular layers. Conversely, if a contact was <1mm above the granular layer, but did not show spiking activity, and contacts above equally did not show spiking activity, all those contacts were considered to be outside the cortex. Similarly, if a layer was >0.75mm below the granular layer, but still showed clear spiking activity (not axon based), we still assigned it to be in the infragranular layers. Based on these approaches, channels outside the estimated range of gray matter of V1 or V4 were excluded from analysis. Note, that layer assignments were made by ^29^, and for consistency we used these assignments (contact by contact) for the data presented here.

### Receptive field mapping

Receptive field (RF) mapping was carried out by a reverse correlation method described in detail previously ^43^. In brief, while monkeys fixated on a fixation point, the RF location and size of neurons in a cortical column was identified performing reverse correlation of spiking activity relative to the presentation of 0.5-2° black squares stimuli (100ms stimulus on with 100ms interstimulus interval) at pseudorandom locations on a 9 × 12 grid (5-25 repetitions for each location) on a grey background.

Offline RFs were determined for each channel via reverse correlation of the MUAe signal (see above for MUAe computation) to stimuli (0.5 – 2 ° black squares). The stimulus-response map was converted to z-scores, after which the RF for each channel was indicated by a contour (thresholded at a z-score of 3) surrounding the peak activity. RF eccentricity ranged from 3.4 - 7.5° in V1, and from 2.5 to 8.9° in V4, and were located in the lower left quadrant. As stated previously, positioning of stimuli in the attention task was based on V1 RFs. The rationale was that these were usually quite a bit smaller than V4 RFs, whereby overlap of V1 and V4 RFs would ensure that V4 RFs also get activated, while the opposite might not be true, if stimuli were centered on V4 RFs. In most cases overlap between V1 and V4 RFs was >80% see figure 1C in ^11^.

### Population coupling computation

To calculate the PC of a neuron, we first computed the population rate of neighbouring neurons (those recorded at different electrode channels, i.e. always excluding the activity from the channel the single unit was recorded from). The population rate was calculated by accumulating the single-unit and multi-unit activity of each included channel (electrode contact) at 1ms resolution. The population rate was then smoothed with a Gaussian kernel of half-width 12 *ms*, followed by subtracting the mean of the smoothed population rate. Then, for every single neuron, its firing rate was calculated within 1ms bins, smoothed with a Gaussian of half width of 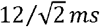 followed by mean subtraction. Finally, we calculated the cross correlation of the mean subtracted smoothed spike train of each neuron with its mean subtracted smoothed population rate. The cross correlation of the smoothed spike train of a neuron with its smoothed neighbouring accumulated spikes was defined by:

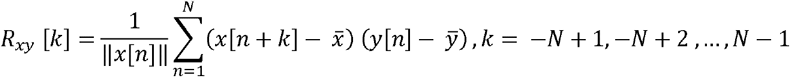

Here, *x* [*n*] represents the smoothed spike train of a neuron, *y* [*n*] represents the smoothed population rate, ∥ *x* [*n*] ∥ represents the norm of *x* [*n*] (that is, the number of spikes fired), 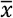 represents the baseline level of the smoothed spike train of a neuron, 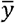 represents the baseline level of the smoothed population rate, k represents the lag of cross correlation, and is the number of samples of *x* [*n*] or *y* [*n*] (*x* [*n*] and *y* [*n*] have the same number of samples). The output vector of the cross correlation was given by:

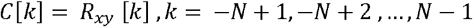

Here, *C* represents the output of the cross correlation and *k* represents the lag of cross correlation. Hence, the population coupling of a neuron at time lag *k* = 0(the time of the lag is 0) was:

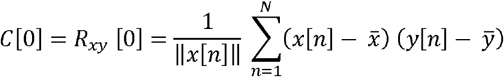

We also normalized the strength of population activity by normalizing the cross correlation of the smoothed SUA and smoothed population activity by their covariance. The normalized degree of population coupling of each SUA was given by:

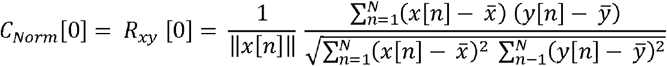

Finally, we calculated the Fisher transform of the correlation coefficient of population coupling (*C*_*Norm*_ [0]) of each SUA as:

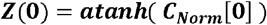

#### Leakage controls

Given the electrode contact spacing had an intercontact distance of 150um it could be argued that some single units might contribute to the multi-unit activity one contact apart (we refer to this as ‘leakage’), which might in turn confound the results. To account for this possibility we recalculated the strength of coupling for all units, when only contacts with distances of 300um and more were used for estimation of population activity.

#### Rate matching controls

Attend RF and attend away conditions can result in different firing rates. To ensure that rate differences did not account for changes in population coupling or other measures, we performed rate matching controls, whereby for each unit we removed trials with the lowest activity (in either condition), until the average activity between the attention condition had less than a 2% activity difference.

#### Response controls

As a further control only neurons with significant stimulus responses were included in the analysis. Here we compared the pre-stimulus response (-250 to 0 ms) to the stimulus onset response period (50-300ms) (Wilcoxon signed rank test), and only selected neurons where the difference was significant.

### Hidden Markov Model and ON-OFF state estimation

ON-OFF state dynamics/periods for this dataset were obtained from the following repositories (G-Node: https://doi.gin.g-node.org/10.12751/g-node.b0mnn2; https://gitlab.com/JvK/cortical-statecoordination). Given that these map 1:1 onto the data set analyzed here, we could determine firing rates of single units during ON as well as during OFF phases, and link their ratio to PC values. For an example of ON-OFF phases in this data set see Figure 1D in ^11^.

However, to make the process of Hidden Markov Model (HMM) fitting to the data set accessible within the context of this manuscript, we repeat the description of the analysis description here. To quantify ON-OFF dynamics in V1 and V4, we fit a Hidden Markov Model (HMM) to the population activity (MUA) across all laminae for additional details see ^11^For the purpose of HMM fitting we extracted population activity by progressively lowering spike extraction thresholds until approximately 100 Hz spiking activity was detected on each channel between fixation onset and the first dimming event. We fit the HMM both to activity from each individual area, following the procedures described by Engel et al. (2016), as well as to the activity from both areas simultaneously.

Our HMM assumes that spike counts on the recorded channels can be well characterized as a doubly-stochastic process, of which the parameters can be accurately estimated ^44^. In this study, spike counts on each channel are assumed to be produced by a Poisson process with different (constant) mean rates during ON or OFF phases of the underlying ‘hidden’ (latent) process *s* common to all channels that we need to infer ^10^. The mean firing rate on each channel *j* in phase *s* is defined by entry 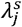 in the emission matrix Λ. The transition matrix *P* gives the probabilities of transitioning between these latent phases. In the transition matrix, each entry indicates the probability of transitioning between two specific phases. For instance, *P*_11_ indicates the probability of transitioning from *s* = 0 to *s* = 0 (remaining in the OFF phase), whereas *P* _12_ indicates the probability of transitioning from *s* = 0 to *s* = 0, more formally: *P*_11_ = *P*_*off*_ = *P* (s_t+1_ = 0 | s = 0 ), *P*_12_ = 1 − *P*_*off*_ = *P* (s_t+1_ = 1 | s = 0 ), These probabilities do not depend on time: at any time step *t*, the probability of transitioning between phases depends only on the value of *s* at time *t* (*s*). The latent dynamics estimated by the HMM thus follow a discrete time series in which *s*_*t*_ summarises all information before time *t*. For each channel, MUA was discretized by determining spike counts in 10 ms bins following each time *t*, with the probability of observing spike count *n* on channel during phase *j* defined as

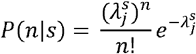

The full description of an HMM is given by the emission matrix Λ, transition matrix *P* and the probabilities π^0^ that indicate the initial values *s*_0_, in which 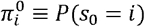 These parameters were estimated using the Expectation Maximization (EM) algorithm ^45^, maximizing the probability of observing the data given the model according to the Baum-Welch algorithm ^44^. Because the EM procedure can converge to a local maximum, rather than the global maximum, we repeated the EM procedure ten times with random parameter initializations and chose the model with the highest likelihood. Random values were drawn from Dirichlet distributions for π^0^ and P, and from a uniform distribution between zero and twice the channel’s mean firing rate for Λ. The EM procedure was terminated if the relative change, computed as | *new* – *original* | / |*original*|, in the log-likelihood was smaller than 10^-3^ and the change in the transition and emission matrix was smaller than 10^-5^, or if it reached the maximum number of iterations (n = 500).

Once the optimal parameters were estimated, we used the Viterbi algorithm to determine the most likely latent trajectory for each individual trial. We applied the HMM separately to each attention condition. For every trial, we applied the HMM during multiple time periods of the task, during fixation and during the time window from 400 ms after cue onset to 30 ms after the first dimming event.

To determine what number of latent phases best described the data, we fit HMMs with the number of phases ranging from 1 to 8, and used a four-fold cross-validation procedure to compute the leave-one-channel-out cross-validation error for each HMM ^10^. We fit the HMM to a randomly selected subset of 3/4 of the trials and computed the cross-validation error on the remaining 1/4 of trials. This procedure was repeated 4 times using a different 3/4 of trials for training and 1/4 of trials for testing the HMM. We computed the cross-validation error *CV*_*var*_ for each channel *j* across all trials *K* and time bins *T* as the difference between the actual and expected spike count according to:

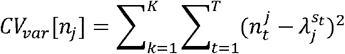

We normalized *CV*_*var*_ to the error in the 1-phase HMM, averaged across channels, cross-validations and conditions, and determined the difference in *CV*_*var*_ with each additional phase in the HMM. For most recordings, and for both V1 and V4, *CV*_*var*_decreased with the addition of a second phase but did not decrease much further with additional phases. This allowed the identification of the elbow (kink) in this error plot as the model with two phases. We included areas/recordings for further analysis that revealed a reduction in cross-validation error of at least 10% with the addition of a second phase but did not decrease by more than 10% with additional phases. For a small subset of recordings, a three or a four-phase model fit the data best [V1: n= 7; V4: n = 1], suggesting that these recordings could contain states with systematically occurring nested fluctuations; these recordings were excluded from further analysis (and are not included in the previously mentioned number of penetrations). In total, we found a reduction of >10% in cross-validation error when fitting a 2-phase versus 1-phase model in 63 V1, and 64 V4 recordings. For these recordings, epoch duration distributions closely followed an exponentially decaying function, consistent with HMM assumptions, indicating that short epoch durations were most prevalent.

While the two-state HMM segments the data into discrete ON and OFF phases, our results do not depend on the assumption of discrete phases. Previous work showed that the ON-OFF dynamics can be also modeled with a continuous latent variable, in which case the inferred firing rates showed bimodality and dynamics consistent with that inferred by the HMM ^10^.

### Ratio of firing rate of ON-OFF phases

In each penetration, HMM was fitted to the MUA to extract ON and OFF phases durations in each trial in two different time windows: 1) Monkey1: 200ms to 500ms after fixation onset, Monkey2: 100ms to 400ms after fixation onset, 2) 400ms after cue onset until 30ms after first dimming onset (identical for both monkeys). Next, we divided the average firing of the ON phase by the average firing of the OFF phase. This provides the ON-OFF firing rate ratio for every trial in the two time periods. We averaged the ratios across trials for each attention condition (Attention-in condition and Attention-out condition).

### Attentional modulation

The effect of selective attention on neural activity was computed via an attention modulation index (*attMI*), defined as:

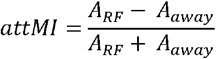

with *A*_*RF*_ as the neural activity when attention was directed towards the RF, and *A*_*awayt*_ the activity when attention was directed away from the RF. This index ranges from 1 to 1, with zero indicating no attentional modulation and with positive (negative) values indicating higher (lower) activity when attention was directed towards the RF. The task included 3 different attention locations, whereby only one overlapped with the receptive field of the neurons under study. For attend away conditions we pooled across the two locations that did not overlap with the neurons’ RFs, as both were equidistant from the attend RF location.

### Noise correlation analysis

We calculated the trial-to-trial spike count correlation (noise correlation) of pairs of neurons in a time window of 500 to 0ms before the first dimming. This was done for attend RF and attend away conditions.

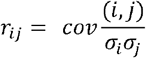

Where cov represents the covariance of the firing rates of neurons i and j, and σ_i_, σ_j_ their respective standard deviation.

