## supplementary figures S1-S7 for "Population coupling of V1 and V4 neurons and its relation to local cortical state fluctuations and attention in macaque monkey"

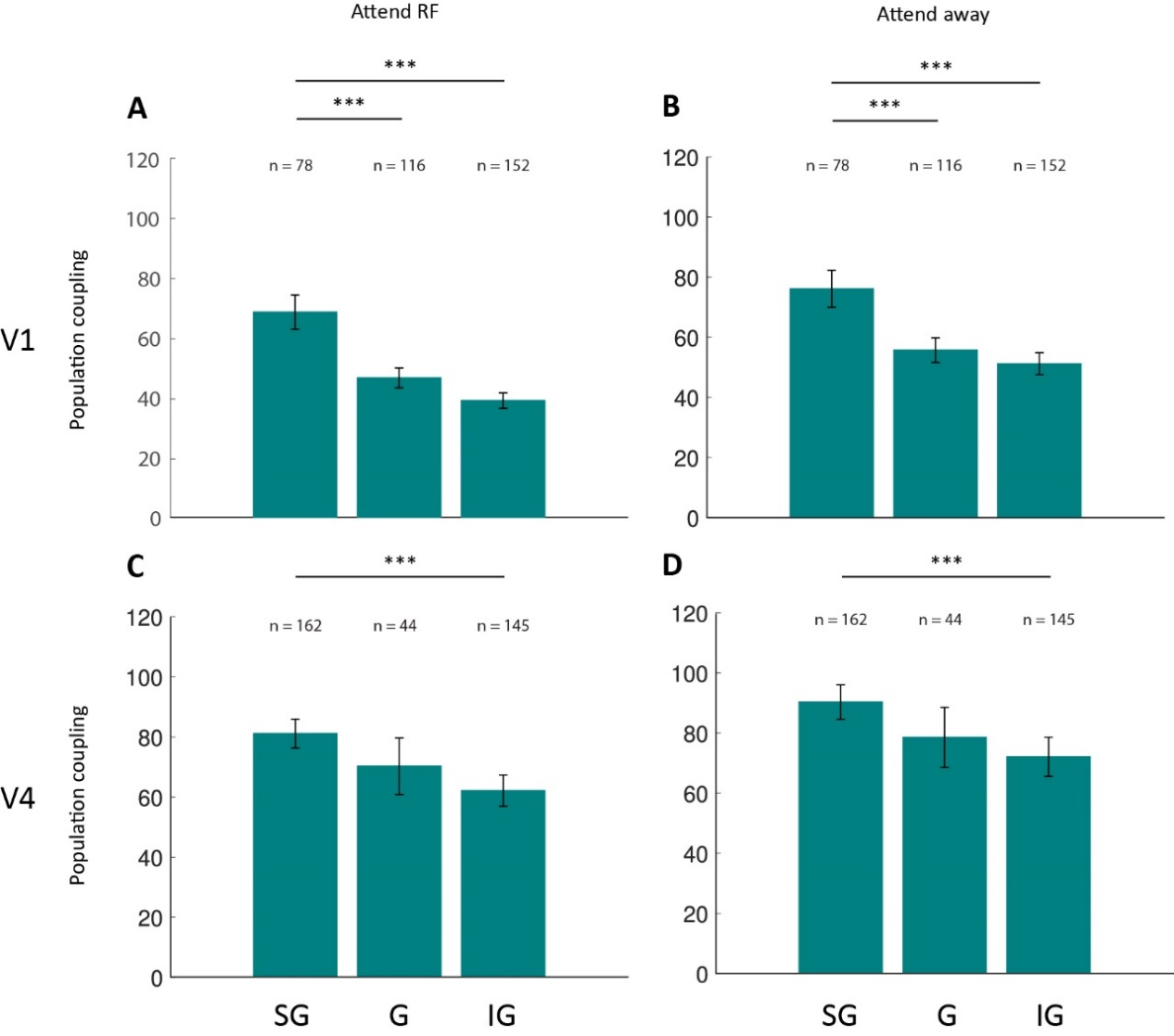

Figure S1: **Population coupling in supragranular (SG), granular (G) and infragranular (IG) layers.** **A)** Population coupling during attend RF stimulus driven activity for neurons located in different layers of V1. **B)** Population coupling during attend away stimulus driven activity for neurons located in different layers of V1. **C)** Same as A, but for V4. **D)** Same as C but for V4. Statistics: Kruskal-Wallis test (Bonferroni corrected); data are represented as means ± SEMs; significance levels *p < 0.05, **p < 0.01, and ***p < 0.001.

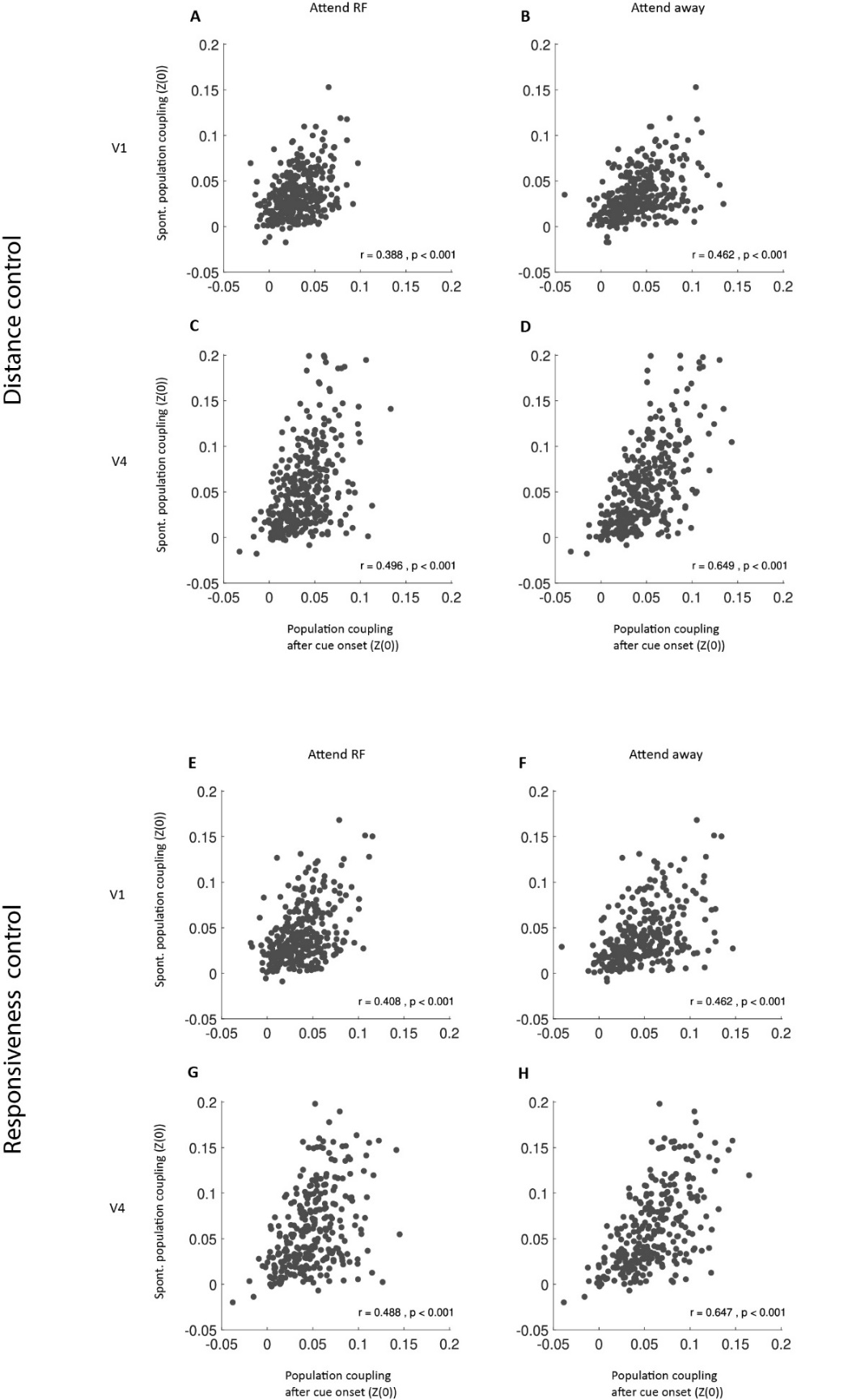

**Figure S2: Population coupling during spontaneous activity compared to stimulus driven activity** (PC quantified by calculating the Fisher transformed correlation coefficient Z(0) between single unit and population activity). **A)** Population coupling during spontaneous vs. attend RF stimulus driven activity for V1 neurons (period after cue onset). **B)** Population coupling during spontaneous activity vs. attend away stimulus driven activity for V1 neurons (period after cue onset). **C)** same as A) but for V4 neurons. **D)** Same as B) but for V4 neurons. **E-H** shows the same comparisons as A-D, but for responsiveness controls. **A-D** shows ‘distance/leakage’ controls (exclu*ding channels less than 200um apart* in population activity calculation). **E-H** shows responsiveness controls (including only single units that responded significantly to stimulus presentations. Insets show correlation coefficient between the two measures, and respective p-values.

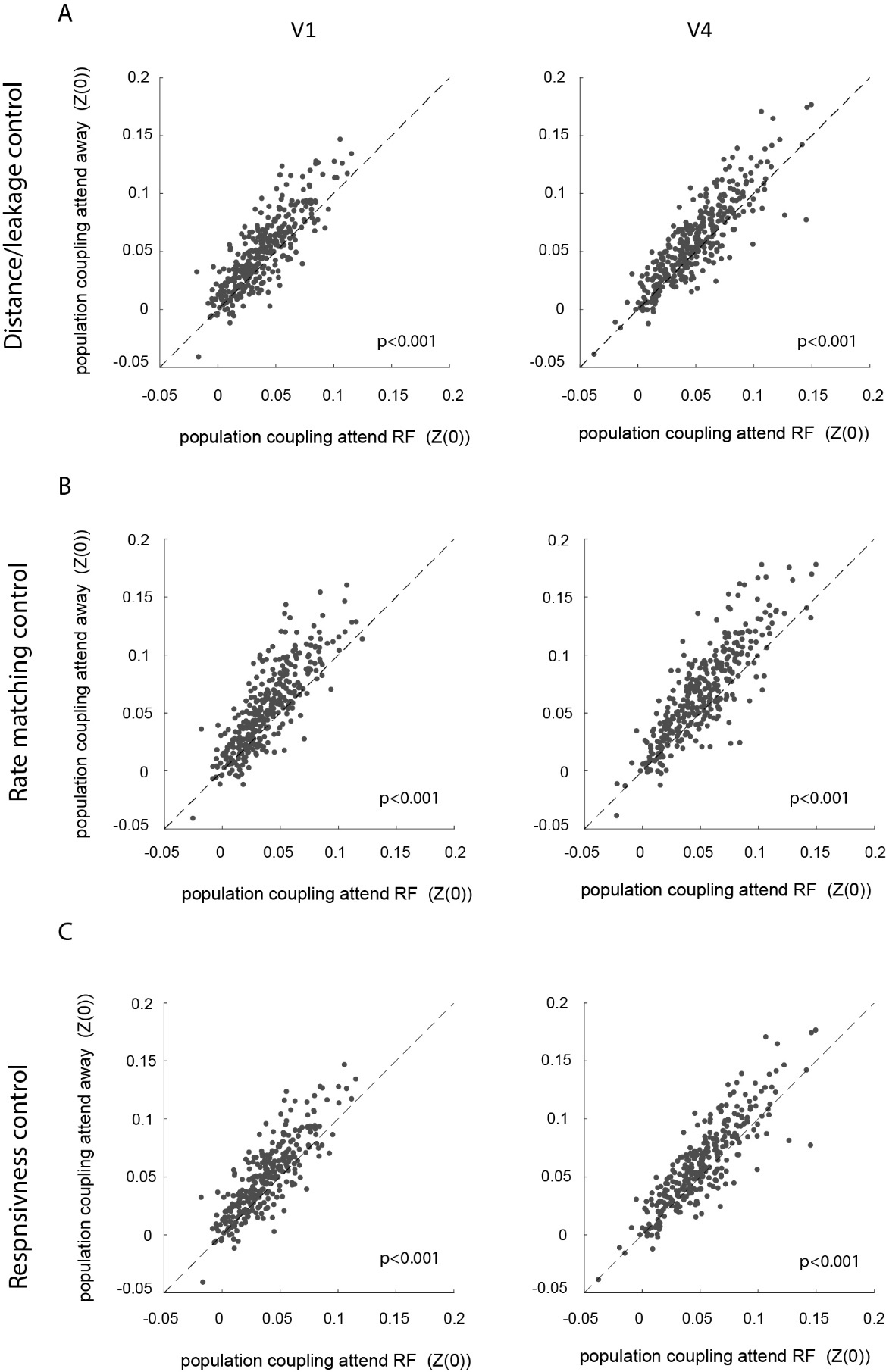

**Figure S3**: **Effect of attention on population coupling of V1 and V4 neurons.** Population coupling (Fisher-transformed correlation coefficient (Z(0)) during attend RF conditions (x-axis) and attend away conditions (y-axis) during the period after cue-onset for V1 neurons (left) and V4 neurons (right). **A**) shows controls for leakage by excluding channels less than 200um apart in population activity calculation. **B**) shows rate matching controls. **C**) shows controls for responsiveness. Insets show p-values of differences in population coupling.

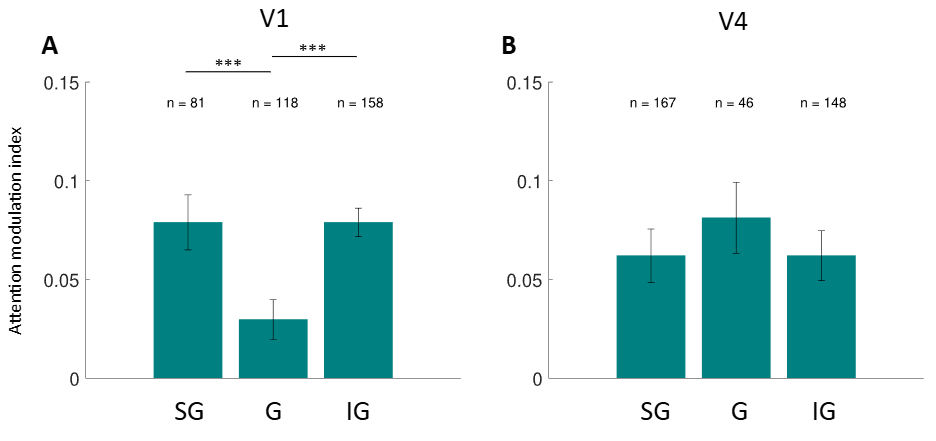

**Figure S4: Attentional modulation indices in supragranular (SG), granular (G) and infragranular (IG) layers. A**) Attention modulation for neurons located in different layers of V1. **B**) Attention modulation for neurons located in different layers of V4. Statistical test: Kruskal-Wallis test (Bonferroni corrected); data are represented as means ± SEMs; significance levels *p < 0.05, **p < 0.01, and ***p < 0.001.

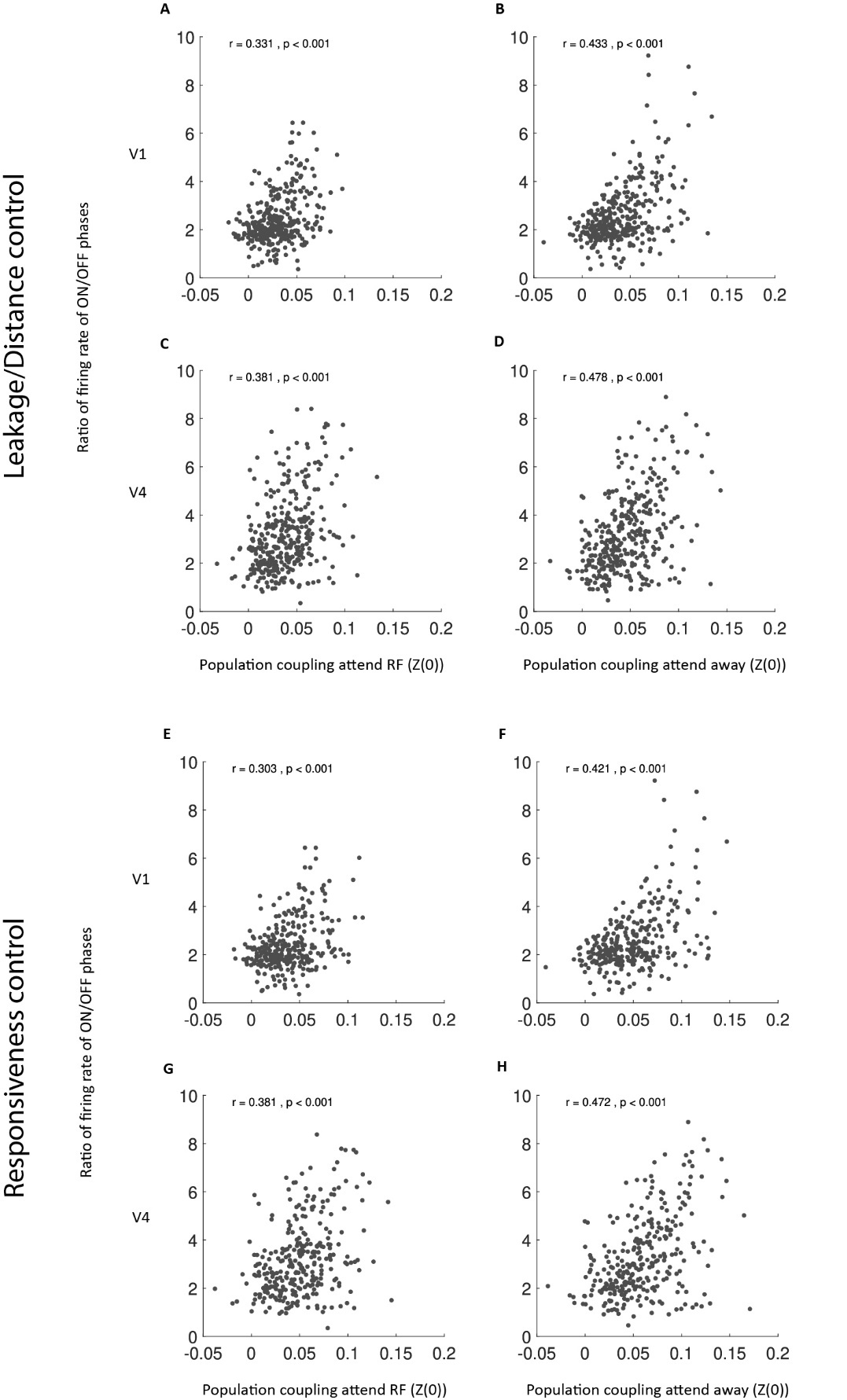

**Figure S5**: **ON-OFF state fluctuation alignment and its relation to population coupling of V1 and V4 neurons.** Population coupling (Fisher-transformed correlation coefficient (Z(0)) during attend RF conditions and attend away conditions during the period after cue-onset for V1 neurons (left) and V4 neurons (right) compared to the ratio of ON-OFF period firing (y-axis). **A-D** shows leakage/distance controls, excluding channels less than 200um apart in population activity calculation. **E-H** shows responsiveness controls, including only neurons that responded significantly to stimulus presentations, Statistic: Wilcoxon signed rank test. Insets show correlation coefficients and p-values.

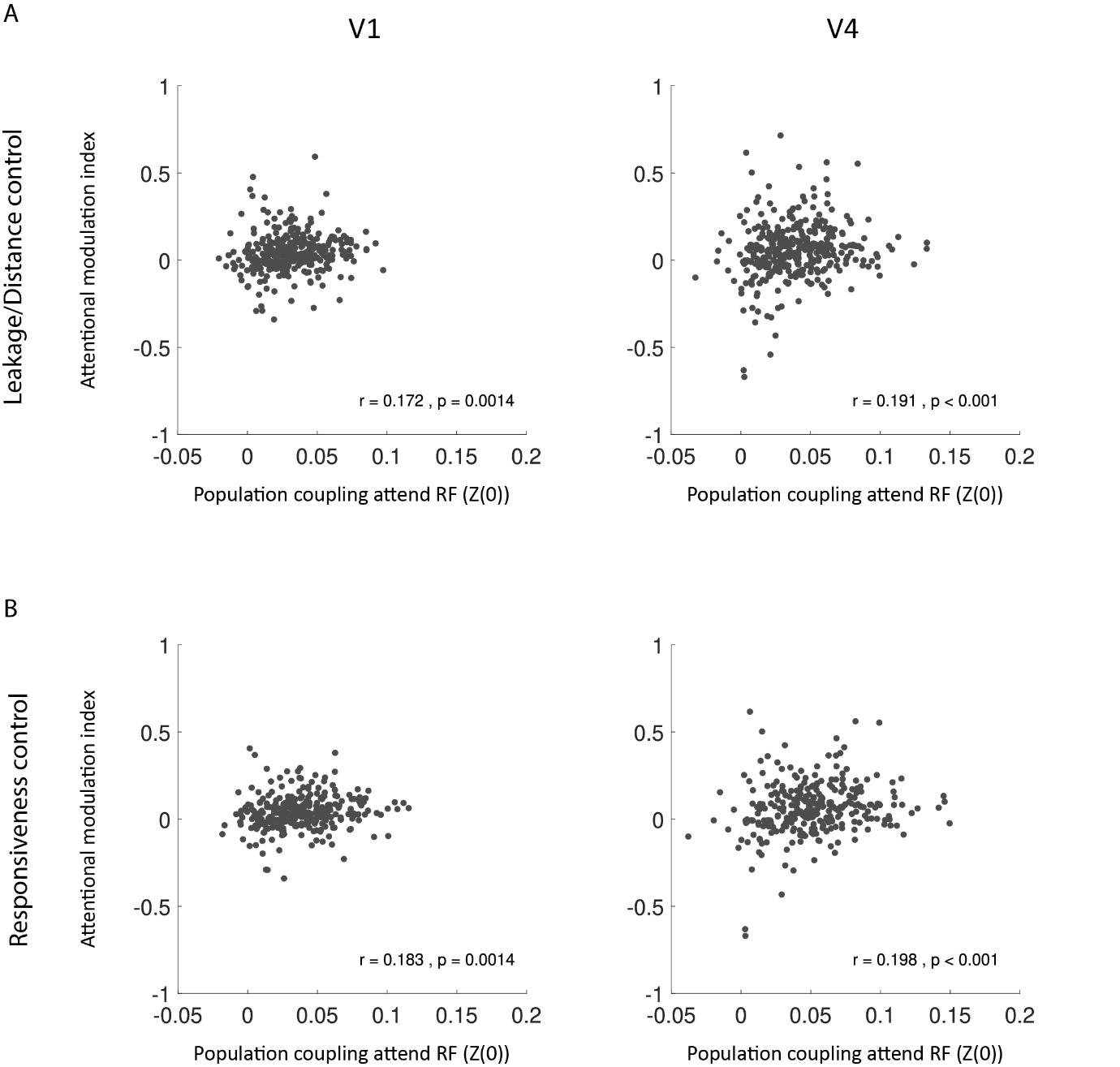

**Figure S6**: **Relation between population coupling and attentional modulation of V1 and V4 neurons.** Population coupling (Fisher-transformed correlation coefficient (Z(0)) during attend RF conditions during the period after cue-onset for V1 neurons (left) and V4 neurons (right) compared to the strength of attentional modulation (y-axis). **A)** shows leakage/distance controls, excluding channels less than 200um apart in population activity calculation. **B)** shows responsiveness controls including only neurons that responded significantly to stimulus presentations, Statistic: Wilcoxon signed rank test. Insets show correlation coefficients and p-values.

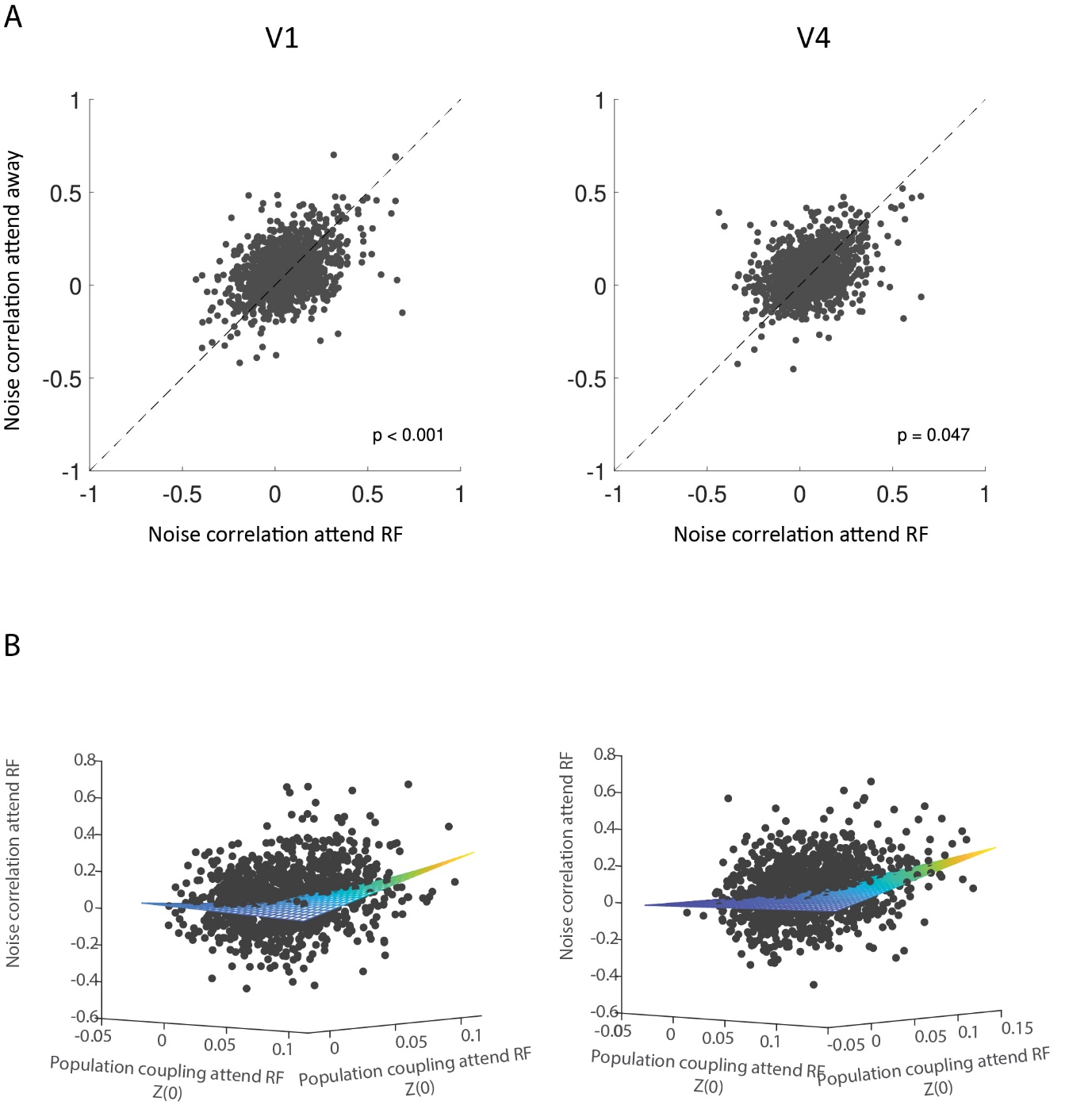

**Figure S7**: **A)** Effect of spatial attention on noise correlation of pairs of V1 (left) and V4 (right) neurons. Noise correlation during attend RF conditions (x-axis) and attend away conditions (y-axis) during the period after cue-onset for pairs of V1 neurons (left) and pairs V4 neurons (right). Insets show p-values of differences in noise correlation. **B**) Relation between noise correlation of pairs of V1 (left) and V4 (right) neurons and population coupling of V1 and V4 neurons during attend RF conditions. Population coupling (Fisher-transformed correlation coefficient Z(0)) during attend RF conditions during the period after cue-onset (x-axis and y axis) for V1 neurons (left) and V4 neurons (right) compared to the noise correlation of pairs of neurons (z-axis). The plane in B) represents the fitted multiple linear regression model predicting noise correlation of pairs of V1 and V4 neurons from population coupling of V1 and V4, including their interaction term (V1: $R^{2}$= 0.053, p<0.001 and V4: $R^{2}$= 0.082, p<0.001).
